# Population Status and Habitat Association of Swayne’s Hartebeest (*Alcelaphus Buselaphus Swaynei* (Sclater, 1892)) in Maze National Park, Southern Ethiopia

**DOI:** 10.1101/2021.01.07.425692

**Authors:** Abraham Tolcha, Simon Shibru, Belayneh Ayechew

## Abstract

We investigated the population status and habitat association of the endemic Swayne’s Hartebeest (*Alcelaphus buselaphus swaynei* (Sclater, 1892)) in Maze National Park, Southern Ethiopia, in 2018 and 2019. Sample count method line-transect was used for the population estimation, while habitat association was made based on the abundance of individuals counted in each habitat. Data were analyzed using descriptive statistics and compared with χ2 test. The total estimated populations of Swayne’s Hartebeest (SHB) in the study period were 1456 and 1492 during wet and dry seasons, respectively showing no seasonal variation. Among the total estimated population, 31% were adult males, 38.46% adult females, 13.97% sub adult males, 15.94% sub adult females and 1.07% young. The number of adult females was higher than the other age groups followed by adult males in both seasons. Significant differences were reported among age and sex structure of population size during both seasons (wet season: χ2= 58.423, df =3, P < 0.05; dry season: χ2=534.079, df= 4, P < 0.05). The maximum group size was 36 and the minimum was 1. The ratio of adult males to adult females was 1:1.24 and 1:1.24, sub-adult males to sub adult females was 1:1.16 and 1:1.12, adult males to sub-adult males was 1:0.36 and 1:0.56, adult females to sub-adult females was 1:0.33 and 1:0.49 in the wet and dry seasons, respectively. The male to female ratio was 1:1.22 and 1:1.19 during wet and dry seasons as well. The population trend among ten years were significantly differed (χ^2^ = 1.708, df= 9, P< 0.05). The SHB was distributed into three types of habitat (riverine forest, open grassland and scattered tree) with significant differences (χ2=1109.937, df = 3, P < 0.05). The savannah grass land was most preferable habitat followed by scattered tree habitat. Maintaining its critical habitat was highly recommended for sustainability of current population status.

## Introduction

Ethiopia is known for a high rate of faunal and floral endemism and diversity, comprising at least 55 endemic mammals, including Swayne’s Hartebeest (*Alcelaphus buselaphus swaynei* (Sclater, 1892)) [1, 2]. Ethiopia’s jagged topography and varied climatic conditions have gifted the country with enormous wildlife species of scenery in Africa [3]. There are eight subspecies of hartebeests, of which Swayne’s Hartebeest is the one [4]. Among these, Ethiopia is home for the three subspecies (i.e. *Alcelaphus buselaphus lelwel, Alcelaphus buselaphus tora* and the endemic *Alcelaphus buselaphus swaynei*) of which all are categorized as endangered. The Swayne’s Hartebeest (SHB) is long-faced, having a rich chocolate brown color of the three sub-species [3] with fine spots and white tips on its hairs. Its face is black save for the chocolate band below the eyes and the shoulders and upper part of the legs are black [5].

The habitats of SHB have been confined by high human settlers and associated livestock populations while, competitions with the cattle for resources (grass) have increased in all the protected areas (PAs) that in turn deteriorated the preferable grass species identified in MzNP and could increase in shrubby and other unpalatable vegetation communities [6]. The intensive agriculture, livestock grazing and human settlement within and around the PAs and the use of intact vegetation remains are mainly considered as a problem through the country including Maze National Park thus could alter its critical habitat and would challenge conservation status of wild animals [3].

Thus, knowledge of the current population size, age and sex structures of the species and its habitat preference would have great value for effective and sustainable conservation effort too.

Therefore, the aim of this study is to determine the current population size, population structure and the habitat association of SHB in the study area.

## Materials and methods

### Study area

The Maze National Park is located at 460 km southwest of Addis Ababa on the way of Wolaita Sodo-Sawla road in SNNPR. The MzNP lies between 06° 3′ to 06^0^ 30′ N latitude and 37° 25′ to 37° 40′ E longitude. Its altitude ranges from 900 to 1200 meter above sea level and covers total area of 202 Km^2^ [7], The Park named after the river Maze, which traverses through its length and rises from southern parts of the surrounding highland and passes through the park from south to north direction and drains into Omo River. Tributaries to river Maze are Lemase, Domba and Zage. Bilbo Hot Spring, which is situated at the southern part of the park, is a natural beauty of hot water gushes out of the ground forming a fountain; and locally and culturally used as a source of cure by different groups of people. The park is surrounded by five districts of Gamo and Gofa zones, namely; Daramalo in the south and southeast, Qucha in the northern part, Qucha Alfa in the northwest, Zala in the Southwest and Kamba in the South (Fig 1).

**Figure 1.**
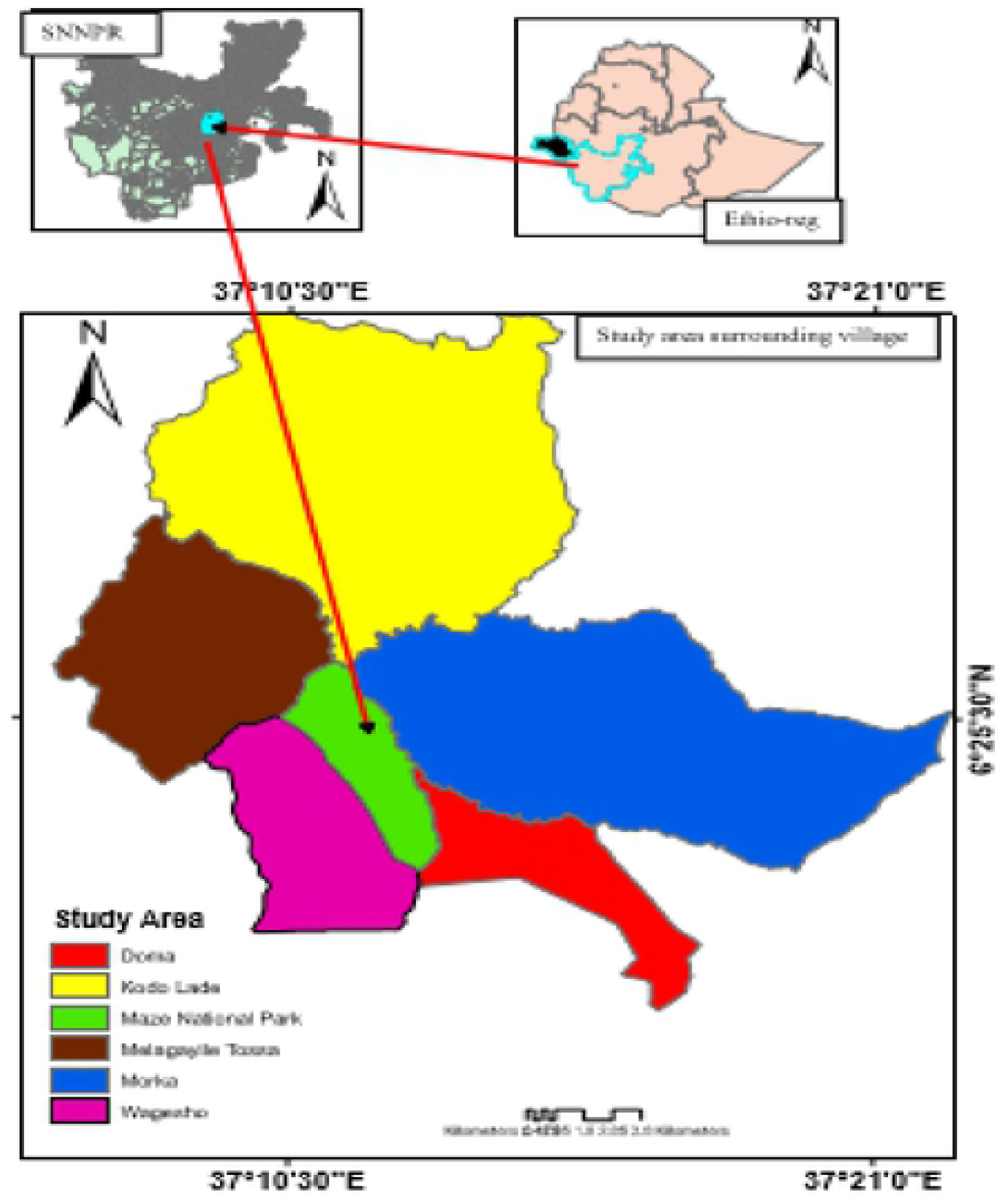
Map of Maze National Park

### Climate

The study area (MzNP) has a bimodal rainfall pattern and typically semi-arid agro-ecological zone of Ethiopia. The annual rainfall ranges between 843 and 1321 mm [8]. Rainy season in Maze extends from March to October, while the dry season is from November to February [3, 8]. The lowest temperature in the wet season is 15.3°C in June and the highest (33.5°C) is in February for the dry season [9].

### Faunal Composition of the Park

About 39 species of large and medium mammals and 196 bird species are found in the Park [10]. The Park is also known for its good population of the critically endangered and endemic sub-species of Swayne’s hartebeest. The existence of these types of resources provides high opportunity for Maze National Park to develop ecotourism. Wild animals are the major natural attractions for ecotourism development [11] such as; Anubus baboon (*Papio anubis*), Vervet monkey (*Cercopithecus aethiops*), Lion (*Panthera leo*), Leopard (*Panthera pardus*), Wildcats (*Felis silvestris*), Serval cats (*Felis serval*), Swayne’s hartebeests (*Alcelaphus buselaphus swaynei*), Klipspringer (*Oreotragus oreotragus*) Waterbuck (*Kobus ellipsiprymnus*), Oribi (*Ourebia ourebi*), Warthogs (*Phacochoerus aethiopicus*), Bush pig (*Potamochoerus porcus*), Reedbuck (*Redunca redunca*), Bush duiker (*Sylvicapra grimmia*), African buffaloe (*Synceros caffer*), Lesser kudu (*Tragelaphus imberbis*) and Bushbuck (*Tragelaphus scriptus*) are among common species recorded in the Park.

### Vegetation

Most of the plains of the MzNP are covered by open *Combretum-Terminalia* wooded grasslands [12, 13]. An occasional variant of woodland vegetation is usually associated with riverine habitats. Combretum dominated wooded grasslands occupy well-drained sites on the upland. This includes the higher ridges and side slopes. It is fire-induced type that replaced a true Combretum woodland or evergreen bush land forest. There are at least 146 plant species were recorded in the Park [13]. Woody plant species like *Combretum adenogonium, Acacia drepanolobium, Maytenus arbutifolia, Harrisonia abyssinica, Acacia seyal, Grewia bicolor, Ziziphus spina-cristi, Bridelia scleroneura, Combretum molle, Pilostigma thonningii* are some of common species in the Park [14] whereas, *Andropogon gayanus, Chrysopogon aucheri, Cyndon dactylon, Dichrostachys cinerea Digitaria abyssinica, Eragrostis cylindrifora, Glycine wightii, Hetropogon contortus, Hyparrhenia hirta, Hyparrhenia rufa, Ischaemum afrum, Loudetia arundinacea, Panicum maximum, Pennisetum thunbergii, Sporobolus panicoides* and *Sporobolus* species, *Themeda triandra, Solanum incanum* are some of grass and herb species are mostly found in the plain area of the Park [15].

### Sampling design

Line-transect sampling methodology was used [16, 17] to collect data. The study area was divided into four different habitat types (open grassland, riverine forest, scattered tree and bush land) based on the major vegetation cover of the study area. Based on the area of the major selected habitat types, ten transects were sampled to cover the major habitat types in the study area. The length of transects was varied from 4.0 to 5.0 km at a distance of 0.5 – 1.5 km between the two nearby transects. Transects were randomly originated and placed with respect to the types of habitat on the map of the study area. Initial points were located using hand-held GPS (GARMIN, etrex 20x). The end point of all transect was found to be reasonably far from their respective habitat edge to avoid the edge effect. Each transect lines were delineated by artificial boundary and natural signs. As a result, from the total potential transects of 27, 11, 9, and 5 in open grass land, riverine forest scattered tree and bush land habitats actual transects of 5, 2, 2 and 1 were randomly selected, respectively (Table 1).

**Table 1.** The number of potential and actual transects in the study area.

| Habitat type | Number of potential transects | Number of sampled transects | Length of each transect (km) | Width of each transect (in km) | The coverage (%) |
| --- | --- | --- | --- | --- | --- |
| Savannah grass land | 27 | 5 | 5 | 1 | 18.5% |
| Riverine forest | 11 | 2 | 5 | 0.5 | 18.1% |
| Scattered tree with grass | 9 | 2 | 4 | 0.5 | 22.2% |
| Bush land | 5 | 1 | 4 | 0.5 | 20.0% |
| Total | 52 | 10 | 18 | 2.5 | 78.8% |

### Data collection

Based on the information gathered during the preliminary survey, survey was conducted on current population status and habitat association of the Swayne’s Hartebeest from October 2018 to April 2019 in Maze National Park including the wet and the dry season.

### Population census

The population status of Swayne’s hartebeest was estimated using the sample survey count method through line transect [16, 18]. Secondary data were used to determine the population trend of Swayne’s Hartebeest for last eight consecutive years since 2010.

During counting, two independent observers were participated to collect the data from the left and the right side of transects in order to increase the validity of the data. Whenever Swayne’s Hartebeest (individual or group) were observed, total number, group size, sex/age group, date, time, altitude, habitat type, and GPS location were recorded [19]. Each habitat type was visited a total of 12 times within a study period. Data were collected twice a day in order to strength the sampling effort; in the early morning (06:30 to 10:30) and late afternoon (14:00 to 18:00) when the animals are active with silent detection [17, 20, 21]. Natural and artificial markings, group or individual size, age and sex composition were taken in order to reduce double counting [22, 19].

### Age and sex structure

Age and sex composition of individual or herd of the animals were recorded as adult male (AM), adult female (AF), sub adult male (SAM), sub adult female (SAF) and young (Yg) [23]. Age and sex determination were carried out based on body size, size and shape of the horn and body color of the Swayne’s Hartebeest [24, 25]. Individuals which are small in their body size were recorded as young and medium in body size were recorded as sub adult male and sub adult female. Individuals those are large in their body size were recorded as adult male and adult female [26].

### Group size

During each sample count, the size of each group of Swayne’s Hartebeest was recorded before classifying into their respective sex and age categories. Animals were considered as members of the same group if the distance between two transects nearby is approximately less than 50 meters, following [27]. Sex ratios for the herds were obtained from direct count of the animals following [28].

### Data analysis

Data were analyzed using SPSS version 20 computer software program and Microsoft excel. Total population was estimated in each habitat following [16]. Number of counted animals during different seasons in each habitat (distribution pattern), age and sex category, herd size was computed using χ2 test. Other data were presented descriptively using tables and figures.

## Results

### Population estimation

The total number of Swayne’s Hartebeest recorded was 1456 and 1492 during the wet and dry season, respectively. The total estimated number of Swayne’s Hartebeest was insignificantly differed between seasons (χ2 = 0.440, df = 1, P > 0.05). Among the total individuals observed in the Park, an average of 31.02% were adult males, 38.53% adult females, 13.95% sub adult males, 15.95% sub adult females and 1.07% were young (Table 2). The number of adult females was relatively higher than the other age groups followed by adult males in both seasons. There was significant difference among age and sex structure of population size during both seasons of the study period (wet season: χ2= 58.423, df =3, P < 0.05; dry season: χ2=534.079, df= 4, P < 0.05).

**Table 2.** Number of individuals in each age and sex categories during wet and dry seasons

| Age and sex structure | Season |  | Mean | Percentage |  |  |
| --- | --- | --- | --- | --- | --- | --- |
|  | Wet | Dry |  | Wet | Dry | Mean |
| Adult male | 481 | 433 | 457 | 33.03% | 29.02% | 31.02% |
| Adult female | 598 | 537 | 567.5 | 41.07% | 35.99% | 38.53% |
| Sub-adult male | 174 | 238 | 206 | 11.95% | 15.95% | 13.95% |
| Sub-adult female | 203 | 268 | 235.5 | 13.94% | 17.96% | 15.95% |
| Young | - | 16 | 8 | - | 1.07% | - |
| Total | 1456 | 1492 | 1474 | 100% | 100% | 100% |

**Table 3.** Age and sex ratio of Swayne’s Hartebeest between seasons of wet and dry.

| Season | Age and Sex Ratio |  |  |  |  |
| --- | --- | --- | --- | --- | --- |
|  | AM:AF | SAM:SAF | M:F | AM:SAM | AF:SAF |
| Wet | 1:1.24 | 1:1.16 | 1:1.22 | 1: 0.36 | 1:0.33 |
| Dry | 1:1.24 | 1:1.12 | 1:1.19 | 1:0.54 | 1:0.49 |

Note: AM= adult male AF= adult female M= male F= female SAM= sub-adult male SAF= sub-adult female.

### Habitat association

Four habitat types were identified in the park, and the Swayne’s Hartebeests were found to be distributed in three of them; namely savannah grassland (SGL), grassland with scattered trees (GST) and riverine forest (RF). The maximum number of population were recorded in savannah grass land i.e., 79.05% and 70.97% in wet and dry seasons, respectively, while the smallest number recorded in riverine forest (1.2%) during dry season only with no in wet season. None of the Swayne’s hartebeest was recorded in bush land (BL) habitat (Fig 2, Table 4). They were significantly differed among t habitat types in distribution (χ^2^=1109.937, df = 3, P< 0.05).

**Figure 2.**
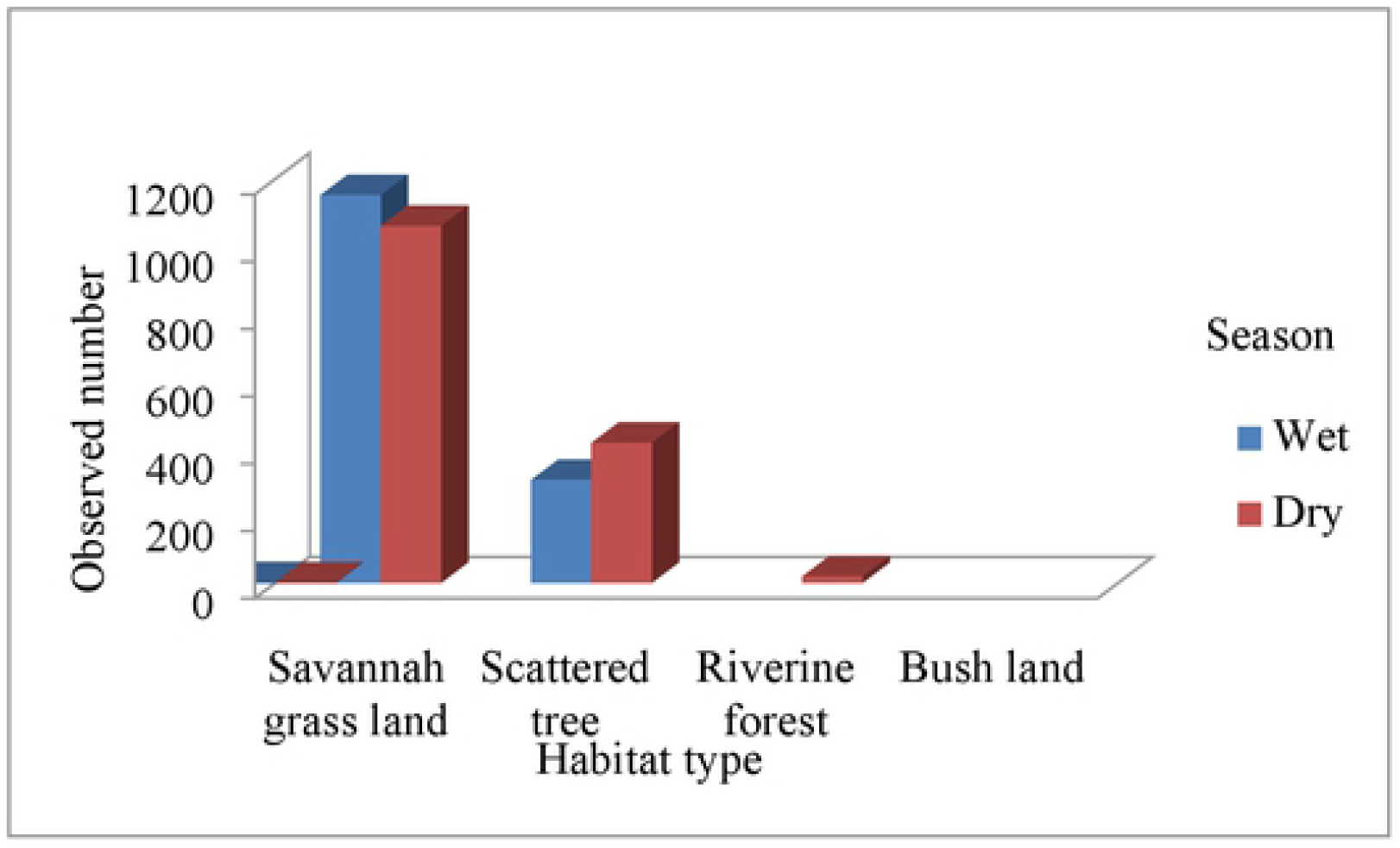
Distribution patterns of Swayne’s Hartebeest with habitat types in seasons

**Table 4.** The number of Swayne’s Hartebeest recorded in different habitat types of wet and dry seasons.

| Season | Type of habitat |  |  |  |  |  |  |  |
| --- | --- | --- | --- | --- | --- | --- | --- | --- |
|  | Savannah<br>grass land | (%) | Scattered<br>tree with<br>grass | (%) | Riverine<br>forest | (%) | Bush<br>land | Total |
| Wet | 1151 | 79.05% | 305 | 20.94% | - | - | - | 1456 |
| Dry | 1059 | 70.97% | 415 | 27.81% | 18 | 1.2% | - | 1492 |
| Mean | 1105 | 75% | 360 | 24.4% | 9 | 1.2% | - | 1474 |

### Group size

The group size also differed with habitat type. The maximum number of individuals in a group was recorded in savannah grass land followed by scattered tree, while the least was recorded in riverine forest i.e. 36, 17 and 23, 25, 5 in wet and dry season, respectively. On the other hand, the minimum group size that revealed in all types of habitat was the solitary male. The group size was differed significantly with habitat types in dry season (χ^2^ = 13.736, df = 1, P < 0.05) and insignificantly differed in wet season (χ^2^ = 5.453, df = 1, P > 0.05). On other hand, significant differences were revealed between maximum and minimum group size between seasons within each habitat type; Savannah grass land (wet: χ^2^ =33.108, df = 1, P < 0.05; dry: χ^2^ = 20.167, df = 1, P < 0.05) Scattered tree habitat (wet: χ^2^ = 14.222, df = 1, P < 0.05; dry: χ^2^ =22.154, df = 1, P< 0.05) whereas, in riverine forest small number was recorded only in dry season.

### Population trend

An increasing trend of population was showed among all study years (Table 5).

**Table 5.** Records of population estimates of Swayne’s Hartebeest among different years in Maze National Park since 2010

| Year | Mean<br>Population<br>size | Trend | References /credit |
| --- | --- | --- | --- |
| 2010 | 371 | Commencement | MzNP office report |
| 2011 | 372 | Stable | Wondimagegnehu and Afework (2011) |
| 2012 | 364 | Decreasing | Yosef *et al.* (2012) |
| 2013 | 614 | Increasing | MzNP annual office report |
| 2014 | 657 | Increasing | MzNP annual office report |
| 2015 | 893 | Increasing | MzNP annual office report |
| 2016 | 998 | Increasing | MzNP annual office report |
| 2017 | 1105 | Increasing | MzNP annual office report |
| 2018 | 1211 | Increasing | MzNP annual office report |
| 2019 | 1474 | Increasing | Present study |

Significant difference was observed among ten years of population trend ( $\chi^2 = 1.708$ , $df = 9$ , $P < 0.05$ )

## Discussion

### Population trend

Swayne’s Hartebeest was locally extinct in some of the country’s National Parks like in Awash National Park [3] and in the Nech Sar National Park [29] while in the Maze National Park the population of SHB was increasing for last ten consecutive years since 2010, wth a slight fluctuation (Table 5). These could be due to reduced human encroachment on the Park. Despite the increasing number of livestock population and related competitions for resources, the Maze national park was still having a potential of good conditions to carry different wild animals particularly the endemic and endangered species of Swayne’s Hartebeest. This was agreed with other studies in Ethiopian protected areas including the present study area [3].

### Group size

The highest group size of Swayne’s Hartebeest was recorded in the savannah grass land during the study period; this might be due to an open access of visibility and having its quality forage, thus most of individuals can be gathered into it and make possibility for effective counting effort. A similar phenomenon was reported earlier [30] that group sizes of large herbivores are mostly affected by habitat structure and population density of animals. Among the total estimation of adult male recorded during the study period, 141 (30.85%) are solitary male while female solitary was not observed during the present study. A similar phenomenon was reported in Maze National Park, where most of solitary SHB recorded was male, while female solitary reported in small number by the author [9]. In addition, fluctuation in group size with seasons and habitat types might be due to change in habitat quality in seasons because of different environmental factors. It was similarly reported that changes in habitat structure between sites could be determines the differences in abundance of animals among habitats in SSHBS [31].

### Population structure

The present study showed that adult females were higher than adult males with mean population size of 38.53% and 31.02%, respectively (Table 2). This is at odd with other studies, which reported more adult males of SHB as compared to the female consisting 48% of the total population in the Park [3]. Similarly, slightly male-biased with insignificant different was reported [32] in Nech Sar National Park, which accounted for 50% adult males and 40% adult females. On other hand, the result is agreed with works of [9] thus out of the total individuals observed, 24.5% were adult males and 34.1% were adult females at Maze National Park. Moreover, high number of females was recorded than males in present study. Studies made at Senkelle were in line with the present study [33, 24, 34]. Decreasing in number of adult males rather than adult females might be due to most of adult males were solitary and involving in territoriality, this exposes them for predation and other human induced factors. As the present study showed that young’s were recorded with small number (1.07%) only in dry season, while did not recorded in wet season. This might be due to dry season is the breeding season for Swayne’s Hartebeest. In addition, a slight increase in population size of Swayne’s Hartebeest in dry season in present study might be due to better visibility than wet, helps for effective sampling and breeding season of them in which new born calf were incorporated. Thus, their breeding season can be determined by habitat type and structure. The same trend was reported by [34] that months of peak lactation match with the most favorable period of the year associated with the grass land structure and which is ranges between December and February [35].

### Habitat association

The availability of quality forage and other resources determine the habitat preference and association of ungulates. Similar phenomenon was reported by researcher that habitat requirements of buffalo were closely associated with the availability of surface water, nutritionally rich food and protection in Chebera Churchura National Park [36]. The present study revealed the same trend that Swayne’s Hartebeest was highly associated with savannah grass land, particularly on the newly emerged grass. This might be due to the abundant and quality forage of savannah habitat that was observed during the study period in the Park. This was in line with other studies, thus savannas are known with its high grass biomass and mosaics, preferring areas access to grass, water and cover [37]. It was an evident that increasing number of Swayne’s Hartebeest recorded in scattered tree during dry season than wet season was due to need for green grass under tree shades. On the other hand, very small individuals were recorded in riverine habitat was because of Swayne’s Hartebeest is known for its dry tolerant and water independent animal. Similarly, it was stated that Hartebeests are well adapted to hot and dry climate and relatively independent of water, and survived in the absence of water source in Senkelle Swayne’s Hartebeest Sanctuary [38]. In line with this, seasonal changes in habitat association of mammals could be forced with their essential food and water requirements [36]. In addition, it was revealed that most of environmental influences, such as human activities, un-prescribed fire often occurring in the Park and livestock grazing, determine changes in habitat association with seasons of the Swayne’s Hartebeest. Similarly, it was reported by researchers that a combination of ecological factors including bush fire and livestock grazing considered as factors that the distribution pattern of wild animals in their natural habitats [39].

## Conclusion

The present study showed that the age and sex structure of total population was dominated by more adult females; this indicates good opportunity for breeding success thus knowledge of sex ratio and age distribution of individual animals is an essential for the evaluation of the viability of the particular species. Though there are a number of human encroachment such as livestock grazing, un-prescribed fire, settlement, habitat destruction, the Maze National Park has potential for this flagship species of the Park in particular and endemic of the country in general. Moreover, to maintain the current status and develop keeping with sustainability of Swayne’s Hartebeest and other related large herbivores in the Park, community based conservation and management was highly suggested. The population of Swayne’s Hartebeest revealed an increasing trend for last ten consecutive years including the present study. In addition, the habitat preference of SHB indicated high association with savannah grass land followed by scattered tree with grass; this delivers useful information to design an appropriate conservation strategy for this endemic species and its critical habitat as well.

## Acknowledgements

We would like to thank Arba Minch University for funding to conduct this research. Our grateful thank also goes to all Maze National Park staff for their integrated help, while performing this research work.

